# DeepDisco: A Deep Learning Tool for Rapid Brain Connectivity Estimation

**DOI:** 10.1101/2025.07.10.663897

**Authors:** Anna Matsulevits, Thomas Tourdias, Michel Thiebaut de Schotten

## Abstract

DeepDisco is a cross-platform software that uses deep learning to rapidly predict brain connectivity maps from binary regions or lesions. It offers a user-friendly and scriptable interface, making indirect connectivity analysis accessible for large datasets and scalable for AI frameworks.

---

DeepDisco is a lightweight, cross-platform software tool that harnesses deep learning to infer structural brain connectivity from simple binary lesion or region masks. This solution addresses a major bottleneck in neuroscience: the inaccessibility of complex connectome estimation due to high computational cost, specialized pipelines, or a lack of user-friendly tools. By offering fast and interpretable connectivity predictions across major fiber classes, DeepDisco facilitates broader access to lesion-network mapping and connectomic inference, supporting research in cognition^1^, brain evolution^2,3^, and neurological disorders^4^.

Recent years have seen increasing reliance on network-based brain models to understand cognition, function, and disease^5,6^. Structural connectomes and disconnection patterns have proven informative, enabling identification of mechanisms behind behavior and providing biomarkers for disease^4,7^. However, practical barriers limit the integration of connectivity models into large-scale or clinical pipelines^8,9^. Most methods require resource-intensive tractography or advanced processing steps, often preventing rapid or scalable use. This limitation is especially salient as the field moves toward large lesion datasets and AI-driven modeling frameworks that demand accessible and modular tools^10^.

We present DeepDisco – a deep-learning-based method that estimates disconnection patterns from a brain region or lesion. The tool predicts four connectivity outputs: association (cortico-cortical connections within a hemisphere), commissural (interhemispheric), projection (cortical-subcortical), and whole-brain connectivity. DeepDisco bypasses computationally demanding fiber tracking while retaining biologically informed predictions, enabling users to work efficiently even with large datasets.

DeepDisco is distributed as a graphical interface built in Python, available for Windows, macOS, and Linux. Upon launching the program, the user selects the desired model type (association, commissural, projection, or whole-brain) and inputs a lesion file or folder of files. The interface includes a progress bar and displays the current file being processed. Typical generation time per lesion is under one second. Users may switch model types and rerun as needed. The interface is fully scriptable, allowing seamless integration into automated pipelines, including future AI-based processing systems, foundation models, and population-scale connectomics.

The core of DeepDisco is a 3D U-Net architecture^11^ trained on advanced connectome maps. Training data were generated using a modified version of the Functionnectome framework^12^ to simulate connections from 1,333 pseudo-randomly defined brain regions^13^, resulting in 5,332 distinct connectome maps (4 × 1333 maps). The model was trained using two 3D input channels: a binary region mask and a static average disconnectome. The training was conducted on an NVIDIA Quadro P4000 GPU (8GB VRAM) and optimized using a hybrid loss (mean squared error + mean absolute error), with weight updates via the Adam optimizer and weight decay of 0.0001. Full details, together with model validation and integration into additional frameworks are described in ^14^.

DeepDisco provides an intuitive input-output pipeline. Given a brain region or lesion mask (in MNI152 2mm space), it rapidly outputs a NIfTI map representing predicted connectivity values for the selected fiber category. This output can be visualized or processed with standard neuroimaging tools. The method preserves key spatial and anatomical priors from its training data, delivering connectivity approximations that align with tractography-derived gold standards^15^. Figure 1 shows the user interface and processing flow, while Figure 2 illustrates representative input/output examples for each connection category.

**Figure 1.**
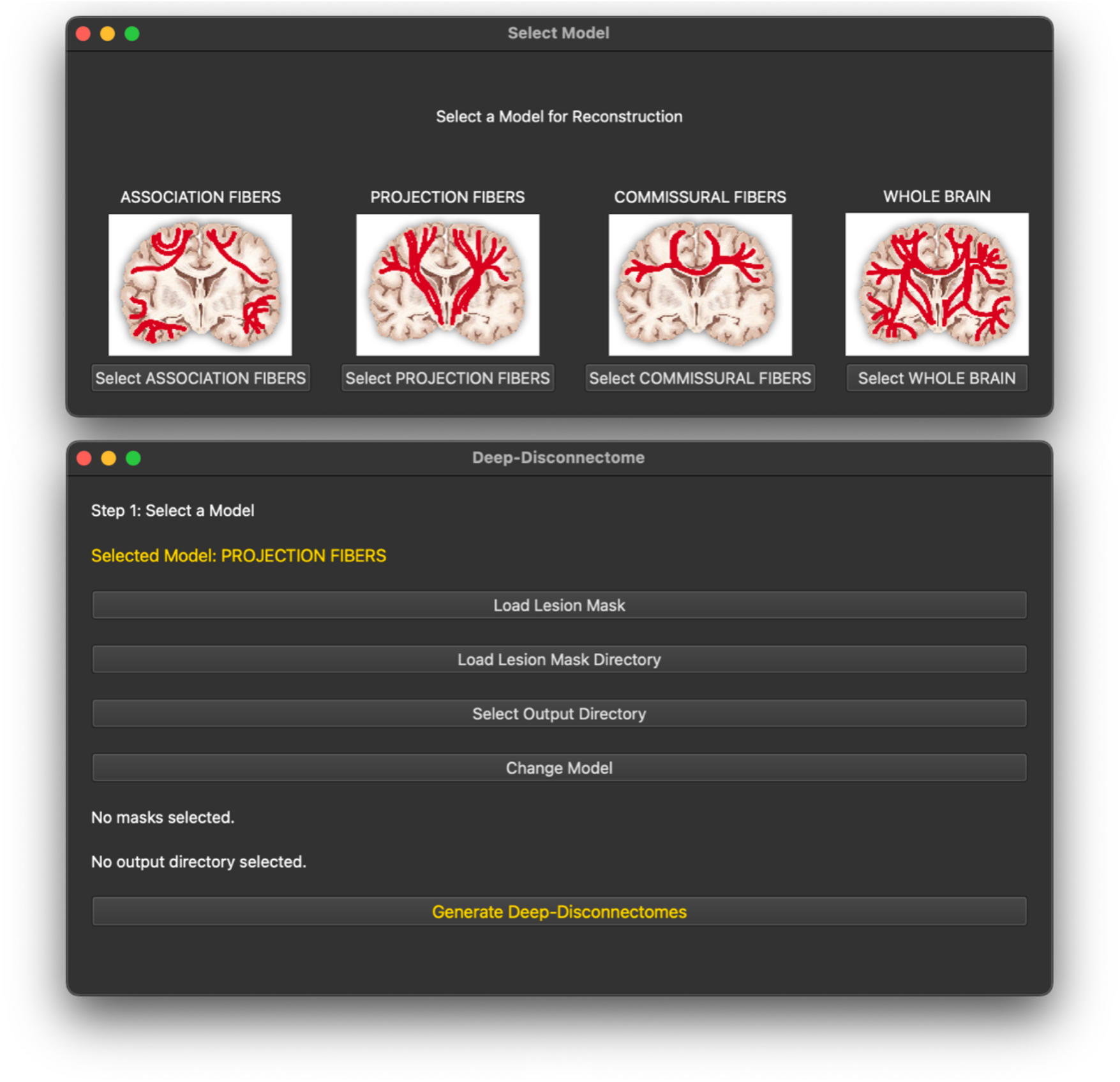
The DeepDisco user interface. The upper window opens upon launching the program and allows the selection of a specified model. The second window is to navigate the interface – loading the input file(s), selecting the output directories, changing the specified model.

**Figure 2.**
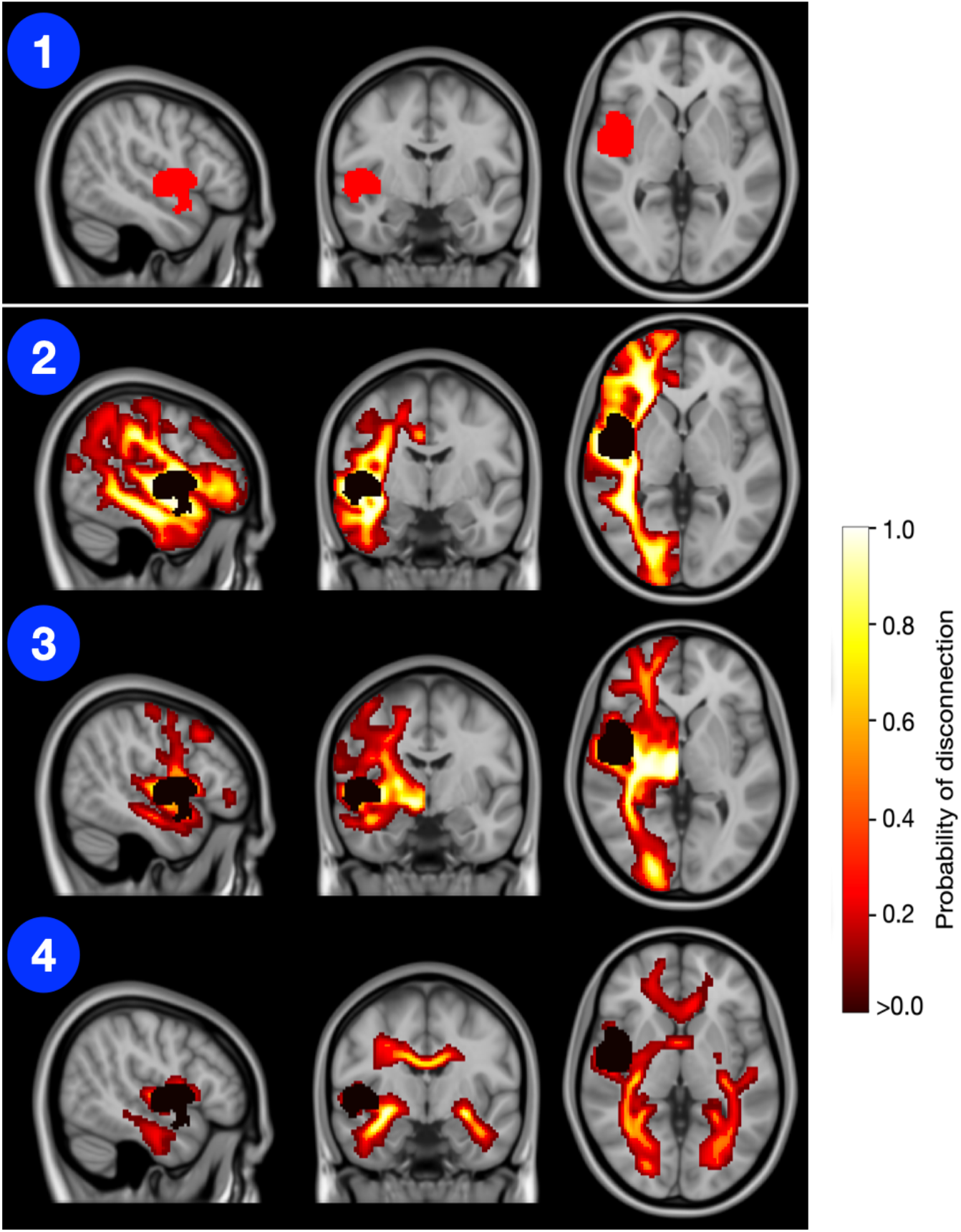
The DeepDisco outputs that can be generated from the binary lesion or region input, marked in red (1). Panel (2) shows the output from the ‘association fibre’ model, panel (3) shows the output from the projection fibre model, and panel (4) shows the output from the commissural fibre model. While the input is binary, the output represents a probability of disconnections, with darker colors (red-black) representing a low probability and lighter colors (yellow-white) representing a high probability for a voxel to be disconnected.

Importantly, DeepDisco is not intended to replace tractography or assess inter-individual variability. Rather, it offers an accessible, validated proxy suitable for large-scale inference, hypothesis generation, or applications where rapid processing is essential.

In conclusion, DeepDisco is a novel software tool enabling rapid, interpretable, and user-friendly estimation of brain circuitry. And, in doing so, allowing the investigation of processes and disorders relying on the interaction between brain regions unavailable in the raw data.

By reducing the computational burden of connectome estimation, DeepDisco opens the door for the wide adoption of disconnection mapping in both research and clinical contexts. The tool is freely available, with detailed documentation and code to support community use and development.

## Acknowledgments

This work was supported by the European Union’s Horizon 2020 research and innovation programme under European Research Council (ERC) Consolidator grant agreement no. 818521 (M.T.d.S., DISCONNECTOME); University of Bordeaux’s IdEx ‘Investments for the Future’ RRI program ‘IMPACT,’ (IMaging for Precision medicine within A Collaborative Translational program) and IHU (Instituts Hospitalo-Universitaires) ‘Precision & Global Vascular Brain Health Institute – VBHI’, ANR-23-IAHU-000, which received financial support from the France 2030 program.

## Code and availability

The DeepDisco codebase is open-source and available at <u>https://github.com/annamatsulevits/DeepDisco</u>.

Binaries and documentation are available via <u>https://github.com/annamatsulevits/DeepDisco/blob/main/README.md</u>.

A user guide and sample data are included.

An online YouTube tutorial is available at <u>https://www.youtube.com/watch?v=CcsmsOFhJtM.</u>

